# Human-like NSG mouse glycoproteins sialylation pattern changes the phenotype of human lymphocytes and sensitivity to HIV-1 infection

**DOI:** 10.1101/404905

**Authors:** Raghubendra Singh Dagur, Amanda Branch Woods, Saumi Mathews, Poonam S. Joshi, Rolen M. Quadros, Donald W. Harms, Yan Cheng, Shana M Miles, Samuel J. Pirruccello, Channabasavaiah B. Gurumurthy, Santhi Gorantla, Larisa Y. Poluektova

## Abstract

**Background:** The use of immunodeficient mice transplanted with human hematopoietic stem cells is an accepted approach to study human-specific infectious diseases, like HIV-1, and to investigate multiple aspects of human immune system development. However, mouse and human are different in sialylation patterns of proteins due to evolutionary mutations of the CMP-N-acetylneuraminic acid hydroxylase (*CMAH*) gene that prevent formation of N-glycolylneuraminic acid from N-acetylneuraminic acid. How changes of mouse glycoproteins chemistry will affect phenotype and function of transplanted human hematopoietic stem cells and mature human immune cells in the course of HIV-1 infection is not known.

**Results:** We mutated mouse *CMAH* on the most widely human cells transplantation strain NOD/scid-IL2Rγ _c_^-/-^ (NSG) mouse background using the CRISPR/Cas9 system. The new strain provides a better environment for human immune cells. Transplantation of human hematopoietic stem cells leads to broad B cells repertoire, higher sensitivity to HIV-1 infection, and enhanced proliferation of transplanted peripheral blood lymphocytes. The mice showed low effects on the clearance of human immunoglobulins and enhanced transduction efficiency of recombinant adeno-associated viral vector rAAV2/DJ8.

**Conclusion:** NSG-*cmah*^*-/-*^ mice expand the mouse models suitable for human cells transplantation and this new model has advantages in generating a human B cell repertoire. This strain is suitable to study different aspects of the human immune system development, might provide advantages in patient-derived tissue and cell transplantation, and could allow studies of viral vectors and infectious agents that are sensitive to human-like sialylation of mouse glycoproteins.

## Background

All vertebrate cell surfaces display a dense glycan layer often terminated with sialic acids that have multiple functions due to their location and diverse modifications [1]. The major sialic acids in most mammalian tissues are N-acetylneuraminic acid (Neu5Ac) and N-glycolylneuraminic acid (Neu5Gc), the latter being derived from Neu5Ac via addition of one oxygen atom by CMP-Neu5Ac hydroxylase (Cmah). The pattern of proteins glycosylation affects the physiology of the cell, cell-to-cell communication, adhesion, migration, recognition by other cells and antibodies [2]. In infectious diseases, sialylation patterns influence how humans interact with some pathogens or viral vectors: including HIV-1, malaria, influenza, and streptococcus [3-13]. In xenotransplantation and stem cell biology, it is a key factor for graft acceptance and preservation of self-renewing properties [14]. Of the two most common sialic acids forms Neu5Gc is widely expressed on most mammalian tissues but has limited accumulation in human cells [15]. The human deficiency of Neu5Gc is explained by an inactivating mutation in the gene encoding CMP-N-acetylneuraminic acid hydroxylase (*CMAH*), the rate-limiting enzyme in generating Neu5Gc in cells of other mammals [16]. This deficiency also results in an excess of the precursor Neu5Ac in humans. This mutation appears universal to modern humans and happens to be one of the first known human-great ape genetic differences with an obvious biochemical readout. In particular, it is important for interaction with Sialic acid–binding Ig-like lectins, or Siglecs. Expression of such lectins vary in their specificity for sialic acid–containing ligands and are mainly expressed by cells of the immune system. For example, humans, compared to mice and rats, express a much larger set of CD33rSiglecs [17]. CD33rSiglecs have immune receptor, tyrosine-based inhibitory motifs, and signal negatively [18]. Interaction with Siglec-7 has the potential to also affect monocyte migration and function [19] along with T-cell activation [8, 20, 21]. During B cells activation and germinal center formation (GC), Siglecs are important for appropriate activation of B cells and responses to T-cell-dependent and independent antigens [22]. In B-cell antigen receptor (BCR) engagement, interaction of CD22 and Siglec-G has been shown to inhibit the BCR signal [23]. Most importantly, exposure to exogenous Neu5Gc is known to cause rapid phosphorylation of beta-catenin in both CMAH-overexpressing cells and bone marrow-derived mesenchymal stem cells, thereby inactivating Wnt/β-catenin signaling and, as a consequence, possibly forcing stem cells to lose pluri- or multipotency [24].

Immunodeficient mice transplanted with human hematopoietic stem/progenitor cells (HSPC) are an established model to study human-specific infections like HIV-1 [25]. However, if a particular sialic acid residue is missing in a donor species (Neu5Gc) and present in the recipient, biologic consequences can be difficult to delineate. Exclusion of mouse Neu5Gc has the potential to improve immune responses to pathogens with non-human patterns of glycosylation like HIV-1 and HCV (hepatitis C virus) and to study the pathogenicity of human-like sialylated pathogen surfaces [12, 17, 26, 27].

To distinguish the effect of the expression of CMP-N-acetylneuraminic acid hydroxylase in mice on human HSPC biology, improve the development of a human immune system in mice, and study responses to HIV-1 infection, we generated mutation in exon 6 of the gene on NSG strain using CRISP/Cas9 technology [28]. We compared original NSG and NSG-*cmah*^*-/-*^ strains for multiple parameters and observed changes in the human lymphocyte phenotype and repertoire. Human lymphocytes generated from HSPC in a human-like sialylation environment exhibit persistence of naïve non-activated T-cell phenotypes and are more sensitive to HIV-1 mediated depletion of CD4+ T-cells. Alternatively, mature human lymphocytes derived from human peripheral blood expand more efficiently in the NSG-*cmah*^-/-^ mice with higher levels of activation.

This new strain expands the utility of the NSG standard strain used to study human hematopoiesis and immunity and allows comparison of new viral vectors for gene therapy and sensitivity to a wider variety of pathogens.

## Results

### Generation of NSG-***cmah*^-/-^ mice.**

To generate a Cmah knockout mouse on NSG background, we designed two single guide RNAs (sgRNAs) targeting exon 6. Schematic of CRISPR targeting are shown in **Fig. 1**. Embryo isolation, microinjection, and generation of founder mice were performed as described in Harms et al [28]. Genotyping of pups showed that one founder contained a shorter sized amplicon. Sequencing of the shorter band revealed a deletion of 27 base-pair sequences in the exon. Genotyping of F1 offspring from this founder is shown in **Fig. 1b, 1c**.

**Fig. 1.**
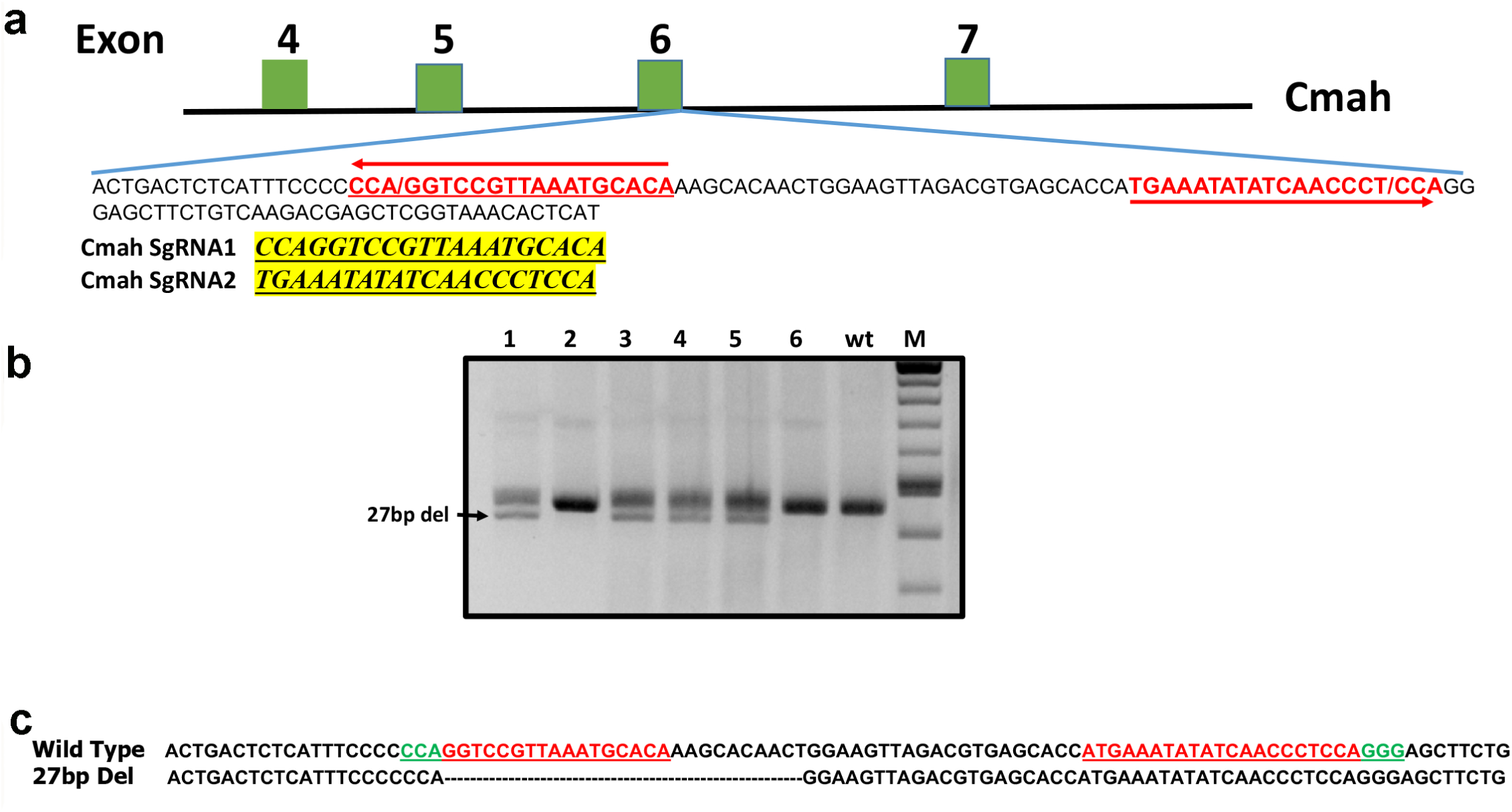
Disruption of Cmah gene in NSG strain mouse using the CRISPR/Cas9 system. **a** Design of 27bp deletion in CMAH gene in exon 6 by sgRNAs. **b** Genotyping PCR of F1 offspring showing wild type and deletion (lines 1, 3-5). **c** Sequence alignment of the wild type and the deletion allele. The guide sequences are shown in red and the protospacer adjacent motif (PAM) sequences are in green.

### NSG-***cmah*^*-/-*^ phenotype**

To confirm the inactivation of *CMAH* gene enzymatic activity and the absence of hydrolysis of Neu5Ac to Neu5Gc, we used the chicken anti-Neu5Gc antibody and anti-chicken immunoglobulin Y (IgY) antibody in different formats: horseradish peroxidase (HRP)-conjugated for Western blot (WB) and immunohistochemistry (IHC) of paraffin-embedded sections (**Fig. 2a and 2b**). FITC-conjugated antibodies were used for analysis of the surface expression Neu5Gc on immune cells in the peripheral blood (**Fig. 2c and 2d**). Neu5Gc expression was undetectable by WB and IHC in all tested tissues: spleen, liver, lung, kidney, heart, gut, and brain. The results were comparable with existing C57Bl/6-Cmah^-/-^ animals. Moreover, flow cytometry showed better reduction of Neu5Gc expression on immune cells of NSG-*cmah*^-/-^ mice than on the existing strain of immune competent animals. By these three techniques, we confirmed the absence of CMAH gene activity and a human-like sialylation pattern of mouse biomolecules. Breeding for two years did not reveal any differences in fertility, body weight, or life span in comparison to the founder NSG mice.

**Fig. 2.**
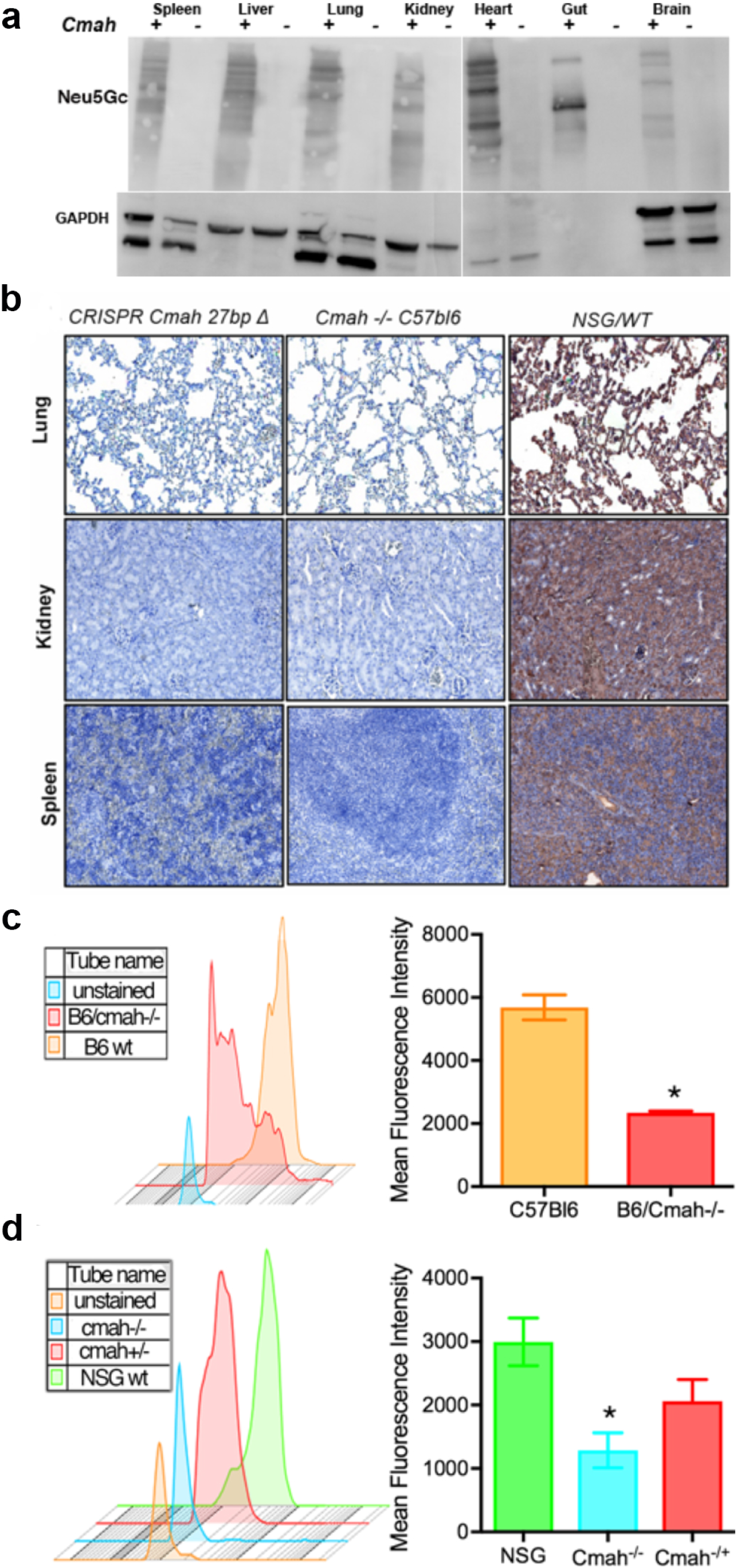
NSG-*cmah*^-/-^ mice phenotype. **a** Western blot analysis of Neu5Gc presence in NSG wild type mice (*cmah*+) and NSG-*cmah*^-/-^ (*cmah*-) tissue samples. All tested mouse tissues with chicken anti-Neu5Gc antibody (Biolegend, CA, USA, 1:4000) were negative. **b** Confirmation of CMAH knockout on NSG background by immunohistology. Spleen, kidney, and lung tissues were fixed, paraffin embedded and 5 μ thick sections of NSG-Cmah^-/-^ generated strain (left), wt NSG (right) mouse and Cmah^-/-^C57/Bl6 original immune competent strain (middle column) were stained with antibodies for Neu5Gc (anti-Neu5Gc antibody, Biolegend, CA, USA, 1:100) at 4 °C overnight. Images captured by Nikon E800 microscope at objective magnification 20×. New generated strain-derived tissues deficient for Neu5Gc as existing *cmah*^-/-^C57/Bl6 strain. Wt NSG mice express Neu5Gc epitopes and tissue sections have brown color. **c** Confirmation of *CMAH* gene knockout on NSG background by FACS. We compared expression of Neu5Gc on white blood cells by staining with anti-Neu5Gc antibody and secondary FITC-labeled anti-chicken reagent. Panel shows Neu5Gc staining for C57Bl/6 *cmah*^-/-^ original strain obtained from the Jackson Laboratories (Stock No: 017588) (red) compare to C57Bl/6 wild type (orange). **d** Panel shows the similar pattern of the absence of Neu5Gc expression on NSG-*cmah*^-/-^ mice (cyan) and the presence of Neu5Gc on cells derived from heterozygous mice that retain enzyme activity and have Neu5Gc on the surface of leucocytes (green and red).

### Comparison of human immune system development after CD34+ hematopoietic stem/progenitor cell transplantation

Another wide application of NSG mice is transplantation of human CD34+ HSPC for the development of human lymphopoiesis [29, 30]. The establishment of human B-cell lymphopoiesis in mouse bone marrow and T-cell lymphocytes in mouse thymus has been well demonstrated [29]. In this model, the expansion of human B cells occurs significantly earlier than CD3+ T cells. We tested whether differences in cell surface glycoproteins sialylation would affect activation of newly generated human lymphocytes. NSG-*cmah*^-/-^ and wild-type (wt) NSG mice were transplanted at birth with the same donor HSPC. At 3 and 6 months post-transplantation, the proportion of human T and B cells as well as activation status was determined (**Fig. 3**). The following markers were used: CD45RA for naïve CD3+ T cells, CD22 for CD19+ B cells. CD22 is a BCR co-receptor that regulates B cell signaling, proliferation, and survival; it is required for T cell-independent antibody responses [31]. The frequency of human T cells in the peripheral blood of 3-month-old mice was higher in NSG wt mice, and the proportion of circulating B cells at the same time was higher in NSG-*cmah*^-/-^ mice. By 6 months of age, there were similar proportions of T and B cells in the peripheral blood (**Fig. 3c**) regardless of sialytion status. In both strains the proportion of CD4+ cells in peripheral blood increased. NSG and NSG-*cmah*^-/-^ mice at this time had similar proportions of naïve CD3+CD45RA+ cells, which declined with time. Human B cells in NSG-*cmah*^-/-^ mice showed a lower number of activated CD19+CD22+ cells at 3 months of age. The differences in B cell activation (CD22 expression levels) were sustained at 6 months post-engraftment. The proportion of CD19+IgD+ B cells was lower in NSG-*cmah*^-/-^ mice at 3 months and did not decline by 6 months as was found in NSG wt humanized mice. The proportion of CD19+IgM+IgD+ B cells in peripheral blood of both strains declined, and the proportion of CD19+IgM^-^IgD+ cells [32, 33] increased. We did not observe significant differences in the numbers of CD19+IgM+IgD^-^ mature B cells between strains at 6 months (5.9 ± 0.9% and 7.7 ± 1.3% NSG-*cmah*^-/-^ and NSG, respectively). The low levels of CD19+IgG+ (0.7 – 3.8%, not shown) cells were found in both strains. The levels of IgM at 6 months of age varied from 1-350 μg/ml (**Fig. 3c**) and only low levels of IgG (1 – 50 μg/ml, not shown) were present in both strains at 6 months of age.

**Fig. 3.**
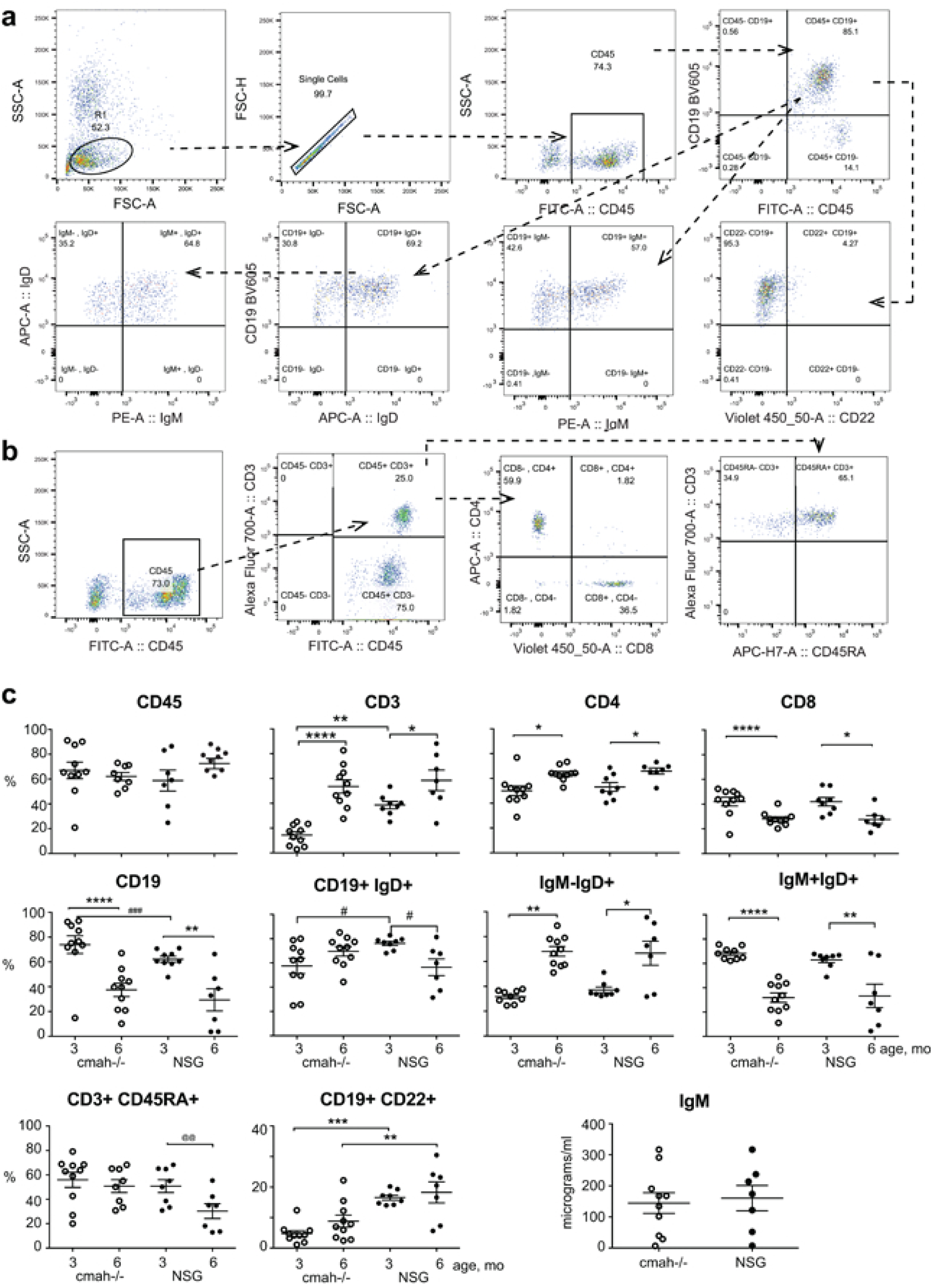
Effects of the *cmah*^-/-^ background on human immune cells expansion and activation after HSPC transplantation. At 3 and 6 months post CD34+ cell transplantation, the frequency of circulating human lymphocyte subsets were analyzed. **a** and **b** Representative plots and gating strategies for human B-cell and T-cell enumeration. **c** Individual mouse NSG-*cmah*^-/-^ (open symbol) and NSG wild type mice (closed symbol) and means with SEM are shown for human B and T cells in mouse peripheral blood. naïve phenotype of CD3+CD45RA+ T-cells exhibited better persistence at 6 months of age in the NSG-*cmah*-/- strain. Lower proportions of mature CD19+IgD+ B cells in peripheral blood of *cmah*^-/-^ mice at 6 months of age, as well as a lower frequency of CD22 B-cell expression at 3 and 6 months of age, were also observed. P values were determined with Kruskal-Wallis test and Dunn’s multiple comparisons tests (*) P values determined by Mann-Whitney test (#) and paired t-test (@) are shown.

### Analysis of T and B cell repertoires in NSG-cmah and wild type NSG mice

To characterize the global B and T cells receptor repertoires, we selected non-fractionated bone marrow cells suspension and spleen tissue samples. Human-specific primers were selected for analysis of human cells according to Adaptive Biotechnologies® (Seattle, WA, USA) technology [34]. We compared the repertoire profiles of bone marrow and spleen within one mouse and between NSG-*cmah*-/- and wt NSG mice. The generation of mature human lymphocytes requires a complex selection process in bone marrow (B cells) and thymus (T cells) and are highly dependent on the microenvironment. Glycosylation of stromal mouse counter-receptors is important for pre-B cells signaling and proliferation [35] and retention in bone marrow for B cells [36]. Maturation and activation of human B and T cells in mouse primary (bone marrow and thymus) and secondary lymphoid organs (spleen, lymph nodes) will likely affect B and T cell receptors repertoire and development [37]. Using the same donor of HSPC, we compared the B and T cell receptors repertoires in the new and existing NSG strain at 8 months of age. Spleen tissue and bone marrow pelleted cells were used for genomic DNA (gDNA) extraction and immunosequencing provided by Adaptive Biotechnologies^®^. Collected data were analyzed using immunoSEQ Analyzer (https://www.adaptivebiotech.com/). We used the common indexes to determine diversity of repertoires based on DNA sequences of the rearranged V-D-J gene segments encoding the third complementarity-determining region (CDR3) of IgH loci and T cell receptor beta chain (TCRβ) in a given sample, the length of CDR3, and usage of IgH and TCRβ genes (**Fig. 4** and additional files 1-5 **Fig. S1 - S5**).

**Fig. 4.**
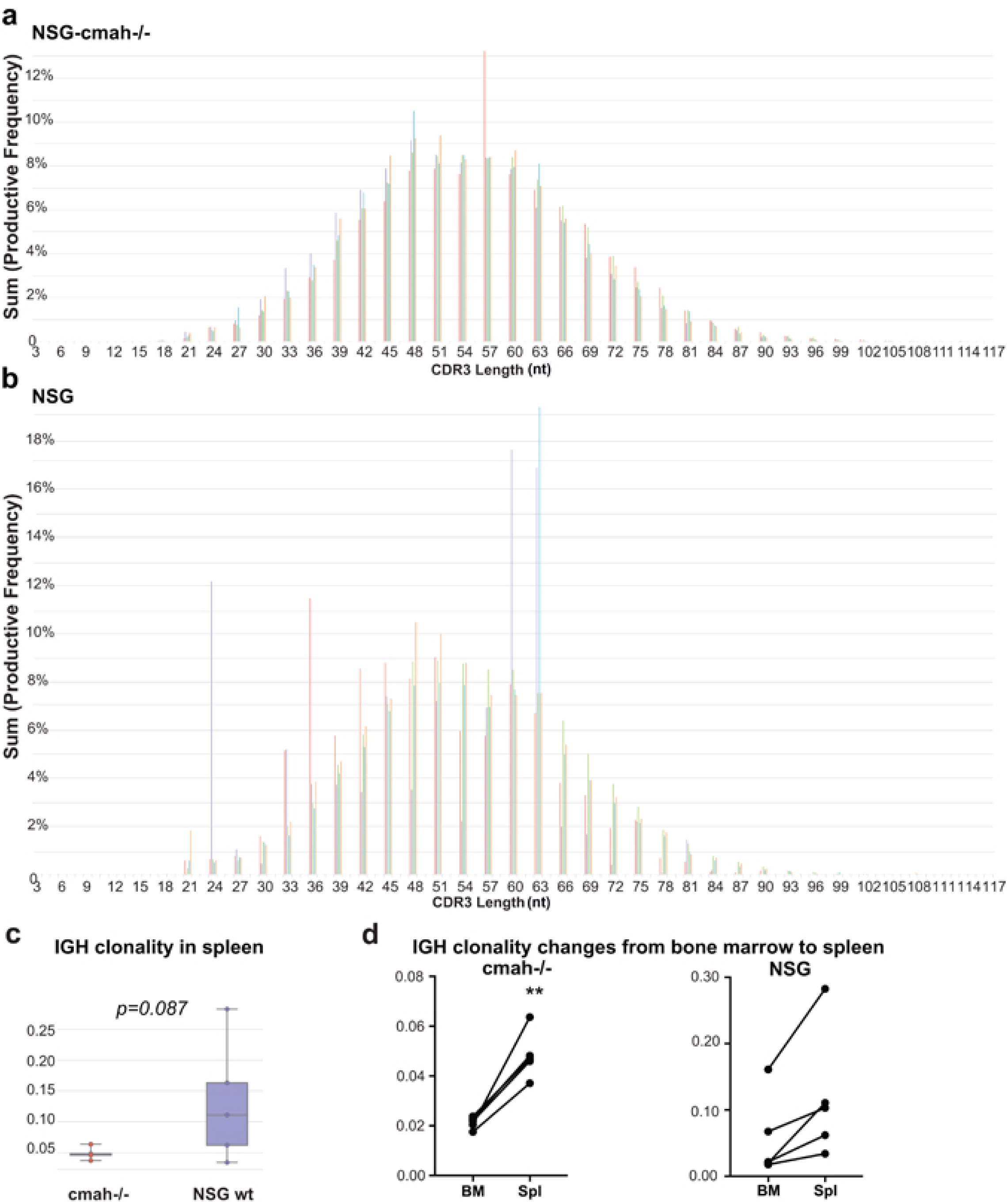
Effects of the NSG-*cmah*^-/-^ background on human immune cells IGH genes repertoires. **a** Histograms of IgH CDR3 length (nucleotides) in spleen of NSG-*cmah*^-/-^ and **b** NSG mice. **c** and NSG-*cmah*^-/-^ background was associated with a lower frequency of clonal B-cell expansion in both bone marrow and spleen. **d** After transition from bone marrow to spleen clonal B-cell expansion increased in NSG-*cmah*^-/-^ mice. Statistical analysis of CDR3 length was performed immunoSEQ Analyzer (https://www.adaptivebiotech.com/). P < 0.05 considered a statistically significant. Five animals per strain were used. Individual CDR3 length in spleen (mature compartment) and bone marrow (developmental compartment) are shown on **Fig. S2.**

Gaussian-like distribution patterns of IgH CDR3 length in NSG-*cmah*^-/-^ mice were similar to wt NSG (**Fig. 4a, 4b**) mice. In contrast, the frequency of clonal B-cell expansion was more common in the wt NSG mice, and the difference was close to statistical significance. A similar size distribution pattern of IgH CDR3 length was observed in bone marrow samples evaluated for five NSG-*cmah*^-/-^ mice. The NSG animals analyzed also had significant variability (Additional file 1, **Fig. S1b**, two animals). The variability in NSG mice was also reflected by variable total number of templates (Additional file, **Fig. S1e**). We observed lower diversity of B cell repertoires in NSG mice compared to NSG-*cmah*^-/-^ strain according to higher clonality of IgH genes in bone marrow and spleen (**Fig. 4c** and additional file 1**, Fig. S1c**). The average IgH CDR3 length naturally generated by the rearrangement machinery was found to be reduced during B cell development and we observed slight shift to the left of CDR3 length in 4 of 5 analyzed NSG-*cmah*^-/-^ mice by comparing bone marrow and spleen compartments (Additional file 2, **Fig. S2**). The use of IgH genes families were similar in spleen and bone marrow (Additional file 3, **Fig. S3C**) and correspond to the known human peripheral blood B cell data [38, 39]. We observed similar changes in IgH D and J gene family usage between bone marrow (pre- and immature B cells) and spleen compartment with mature B cells (Additional file 3, **Fig. S3C**). The clonality of IgH gene loci were two folds higher in spleen compared to bone marrow, and in *cmah*^-/-^ mice reached statistical significance (**Fig. 4d**).

We did not observe significant differences in TCRβ chain gene usage (Additional file 4, **Fig. S4**) nor repertoires in new strain compare to NSG mice (Additional file 5, **Fig. S5**). TCRβ clonality was lower in spleen compared to bone marrow in both strains, but these changes did not reach statistical significance (Additional file 5, **Fig. S5b**). The richness of repertoire was higher in spleen tissues compared to bone marrow samples in both strains (**Fig. 5d**). We did not observe statistically significant differences in TCRβ CDR3 length (Additional file 5, **Fig. S5a, 5b**) in spleen and bone marrow (not shown). Overall CDR3 length (nucleotides) was shorter in T cells compared to B cells. We did not find statistically significant differences in samples clonality and noted higher TCRβ richness in spleen compared to bone marrow as total number of TCRβ chain gene templates. Additionally, we observed better Pielou’s evenness of TCRβ in NSG *cmah*^-/-^ mice and overall increased evenness in spleen compared to bone marrow compartment (Additional file 5, **Fig. S5c, 5d, and 5e**).

**Fig. 5.**
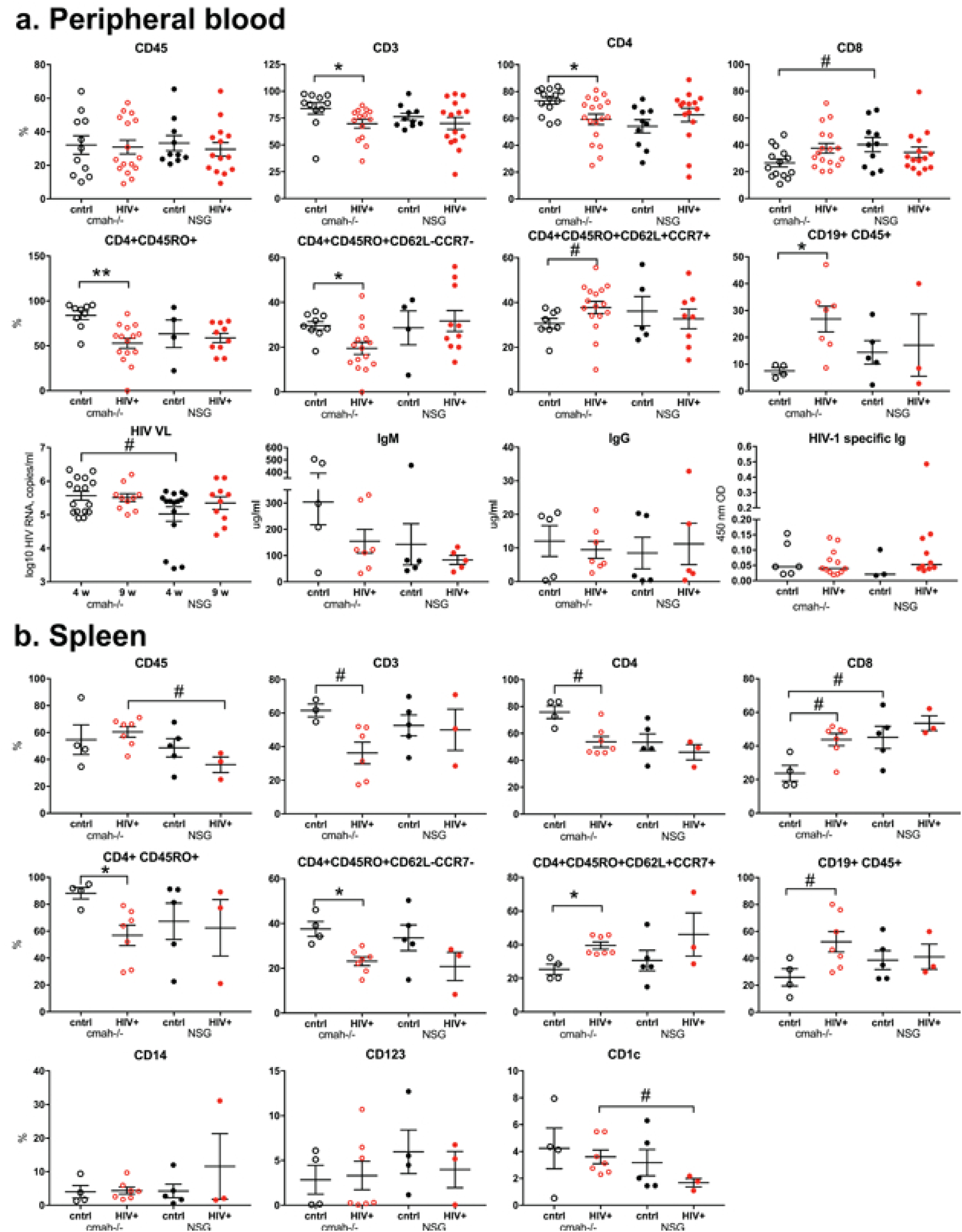
Effects of the *cmah*^-/-^ background on human cell responses to HIV-1 infection. NSG-*cmah*^-/-^ and NSG wt mice were infected with HIV-1_ADA_ intraperitoneally at 6 months of age. **a** At 4 weeks post infection, blood samples were collected for FACS analyses of the peripheral blood. **b** Four to seven animals per group were euthanized for the spleen profile analysis. Bone marrow data shown on **Fig. 6**. FACS gating strategies were used: human CD45/CD3/CD19; CD3/CD4/CD8. For available blood samples, additional analyses for human cells subpopulations CD4/CD45RA/CD45RO/CD62L/CCR7 were done. For spleen, additional analysrs included: human CD45/CD14/CD123/CD1c. Available plasma samples were analyzed for the HIV-1 RNA copies number, human IgM, IgG and HIV-1 specific antibodies at 1:10 times dilution (last panel **a**). Individual mouse and means with SEM are shown. P values were determined with Kruskal-Wallis test and Dunn’s multiple comparisons tests (*) and Mann-Whitney test (#). P ≤ 0.05 were considered significant. Viral load at 4 weeks post infection was compared with unpaired t test with Welch’s correction. In comparison to NSG mice, NSG-*cmah*^-/-^ mice showed a higher sensitivity to HIV-1 infection with increased viral load at 4 weeks post infection and a significant decrease in numbers of CD4+ T cells including effector CD4+CD45RO+CD62L^-^CCCR7^-^ cells post-infection. Profiling results of animals euthanized at 9 weeks post-infection are shown in Supplemental Material (**Fig. S7**).

### Comparison of human immune system responses to HIV-1 infection in NSG-*cmah*^-/-^ and wt NSG mice

NSG mice, humanized by human hematopoietic stem cell transplantation, are a proven model to study the pathogenesis of HIV-1 infection [38]. We evaluated the effects of human-like sialylation of mouse tissues on the sensitivity of human cells to HIV-1. Two groups of mice with similar levels of reconstitution of circulating human CD45+ cells in peripheral blood were infected with the same dose of HIV-1_ADA_ intraperitoneally at the age of 30-32 weeks when the majority of peripheral human cells were CD3+CD45RA+ T cells. Four weeks post-infection, peripheral blood, spleen, and bone marrow samples were analyzed (**Fig. 5 and 6,** Additional files 6 and 7, **Fig. S6, S7**). We observed a significant reduction in human T cells in all three compartments in NSG-*cmah*^-/-^ mice including both CD3+ and CD3+CD4+ T-cells. The total number of CD4+CD45RO+ cells and among this antigen-stimulated cells proportion of CD4+CD45RO+CD62L+CCR7+ effector memory cells, the most sensitive to HIV-1 depletion, decline in *cmah*^-/-^ mice compared to unaffected. The result was an increased frequency of central memory T cells CD4+CD45RO+CD62L+CCR7+ in the peripheral blood and spleen of HIV-1 infected mice. This effect was not observed in NSG mice peripheral blood and spleen compartment. We found a higher HIV-1 viral load (VL) in the peripheral blood of NSG-*cmah*^-/-^ mice at 4 weeks post-infection. In both strains, VL persisted at high levels for up to 9 weeks post-infection. As T cells decreased, we observed an increased frequency of B cells in all three compartments (blood, spleen, bone marrow). We also noted an increased proportion of human CD1c+ cells in spleen of infected NSG-*cmah*^-/-^ mice compared to wt NSG. Corresponding increases in myeloblasts (CD34+CD117+), promonocytes (CD4^dim^CD14^neg^ ^or^ ^dim^), and mature monocytes (CD4^dim^CD14^bright^) in bone marrow of HIV-1 infected NSG-*cmah*^-/-^ mice were observed (**Fig. 6c**). We did not observe any differences in natural killer (NK) and NKT cells subpopulations (not shown) between the two strains. We also did not find differences in the levels of immune globulins and circulating HIV-1 specific binding antibodies. At nine weeks post-infection, in peripheral blood of HIV-1 infected NSG-*cmah*^-/-^ mice, the decreased proportion of CD3+ cells and specifically CD4+CD45RO+ effector memory cells with increased number of monocytes CD14+ were found (Additional file 7, **Fig. S7**).

**Fig. 6.**
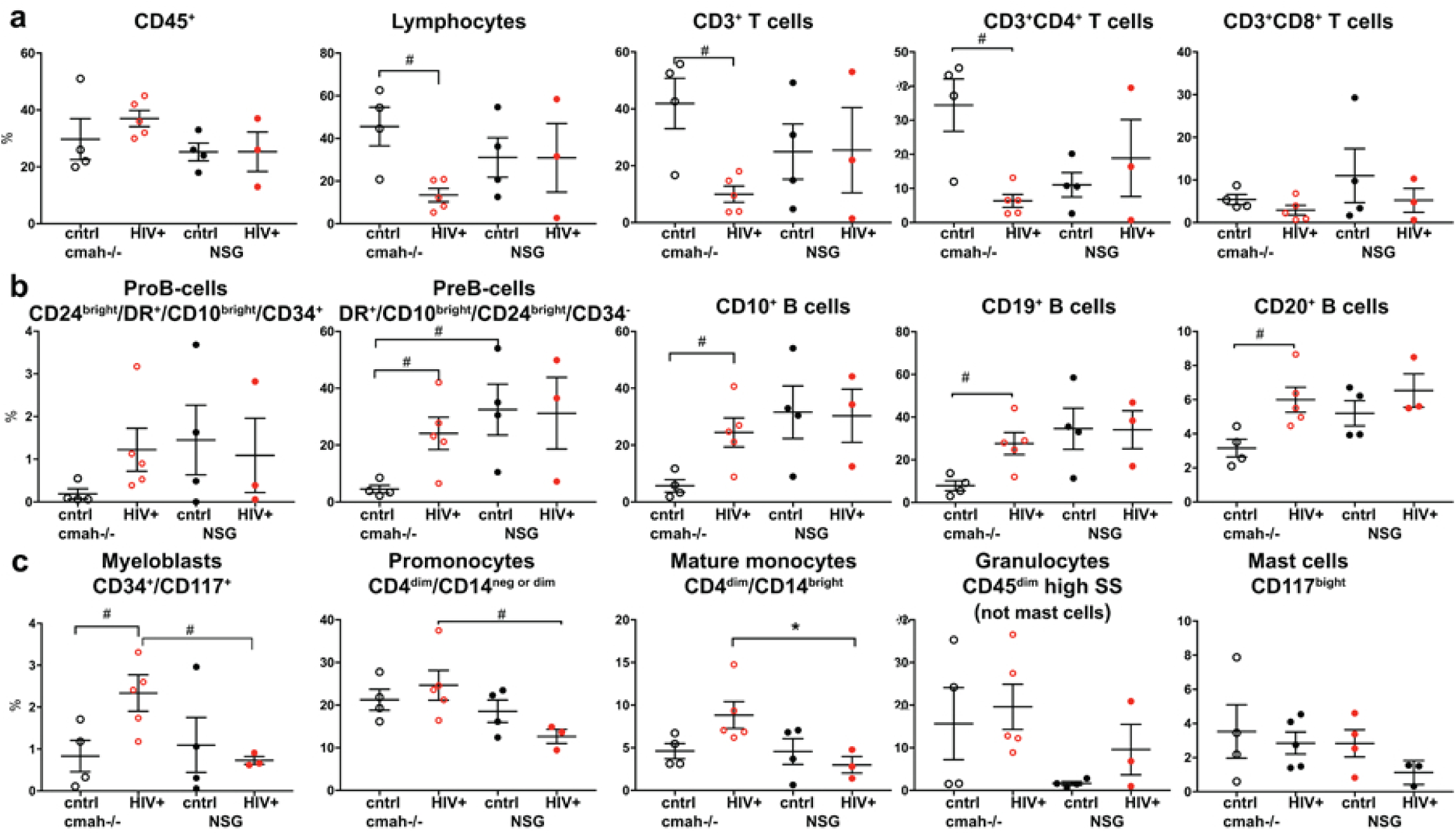
Effects of the *cmah*^-/-^ background on human cell responses to HIV-1 infection in bone marrow. **a** Human differentiated T cells, **b** precursors and mature B cells, **c** myeloid lineage cells FACS analyses. Mice bone marrow were reconstituted with comparable proportions of human CD45+ cells; HIV-1 mediated depletion of T cells were more pronounced in NSG-*cmah*^-/-^ animals compared to NSG wt. HIV-1 infection increased the relative proportion of B cell precursors and mature B-cells (CD20+) and the proportion of CD14+ macrophages compared to NSG mice. Individual mouse and means with SEM are shown. P values were determined with Kruskal-Wallis test and Dunn’s multiple comparisons tests (*) and Mann-Whitney test (#). P ≤ 0.05 were considered significant. The gating strategies are shown in Supplemental Material (**Fig. S6**).

### Effects of NSG-*cmah*^-/-^ phenotype on transplanted human peripheral blood mononuclear cells behavior

NSG mice are widely used for the transplantation of human peripheral blood immune cells (PBMC) [39]. We considered that the absence of Neu5Gc and human-like sialylation of glycoproteins could change the behavior of mature human immune cells derived from the adult donor. PBMC isolated from one donor were transplanted into the NSG and NSG-*cmah*^-/-^ mice intraperitoneally. Starting from day 7 post-transplantation, mouse peripheral blood was collected and stained for the presence of human immune cells (**Fig. 7**). Human T lymphocytes were the major population with low absolute number of B cells and monocytes in peripheral blood. The expansion of human cells in NSG-*cmah*^-/-^ mice significantly (∼2 – 3.5 times) exceeded that seen in the NSG mice, and the absolute count of human CD45+ cells (mean ± SEM) were 16 ± 2 vs 9 ± 1 cells/μl at day 7, 513 ± 135 vs 146 ± 58 cells/μl at day 14, 474 ± 130 vs 168 ± 118 at day 21 (P < 0.05) (**Fig. 7b**). The proportion of human cells in spleen were similar at the end-point of observation (**Fig. 7c**). Human T-cells in the mouse environment became activated and switched expression of CD45RA (naïve) to CD45RO (effector-memory). The activation of CD4+ cells in Cmah^-/-^ mice significantly exceeded the CD45RO to CD45RA switch in wt NSG mice. To a lower extent, the same was observed within CD8+ T-cells (**Fig. 7d**). L-selectin (CD62L) is an adhesion molecule that recognizes sialylated carbohydrate groups, mediates the first steps of leukocyte homing to peripheral lymph nodes, and is programmed for recirculation through lymphoid organs, thus crucially controlling the initiation and maintenance of immune responses to pathogens [40]. CD62L+ T-cells also were generated at higher frequency in Cmah^-/-^ mice, including the splenic population of CD4+CD45RO+CD62L+ (**Fig. 7e**). Overall, for this particular donor, engraftment and expansion of mature human peripheral blood lymphocytes were more pronounced in the NSG-*cmah*^-/-^ mouse strain.

**Fig. 7.**
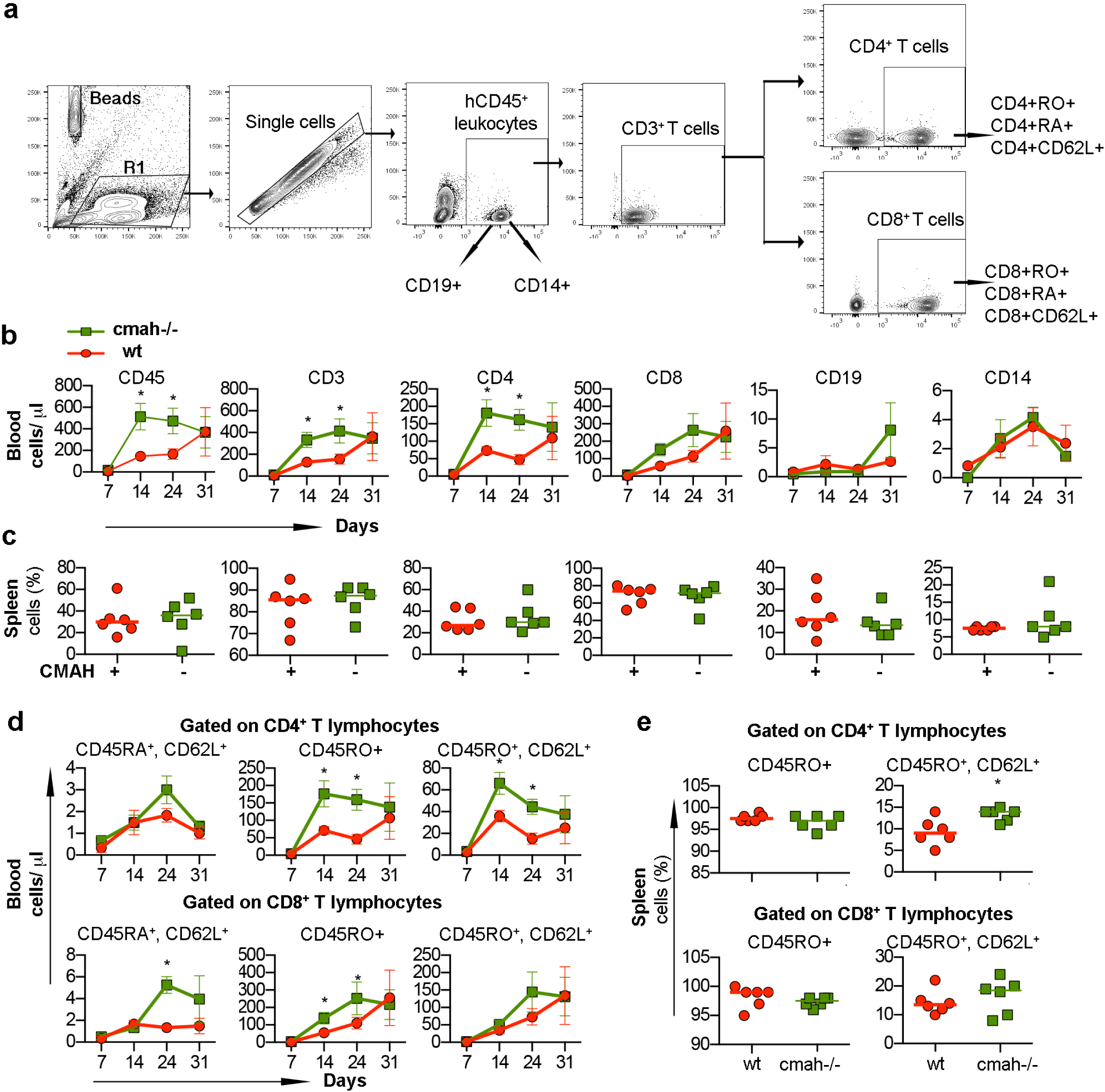
Human immune cells expansion and phenotype changes in NSG-*cmah*^-/-^ mice. Males NSG-*cmah*^-/-^ and wt NSG 5 weeks of age were transplanted intraperitoneally with single-donor human peripheral blood mononuclear cells. The number and phenotype of human cells in peripheral blood were analyzed up to 31 days post transplantation and percentage of human cells in the spleen at the end point of observation. **a** Gating strategy to identify human CD45+ cells, CD19+, CD14+, and CD3+ T cells and their subpopulations. **b** Absolute counts of human cells in peripheral blood with accelerated expansion of human T cells and CD4+ cells in the NSG-*cmah*-/- compared to NSG. **c** Proportion of human cells in the spleens at the end-point were similar in both strains. **d** Proportion of memory CD45RO+ and central memory CD45RO+CD62L+ were increased in NSG-*cmah*^-/-^ mice among CD4+ T-cells in blood and spleen. **e** Proportion of CD8+ T-cells were also increased but to a lesser degree. Six animals per group were used and **b** and **d** represent means with SEM, **c** and **e** individual mouse data. Statistical analysis done by 2-way ANOVA with Sidak’s multiple comparison test, * - P < 0.05.

### Host glycoproteins sialylation pattern affects the clearance of HIV-1 virus, half-life of infused human IgG and rAAV2/DJ8 biodistribution

There is evidence that evolutionary loss via mutation of the CMAH genes changed the sensitivity of humans to viral and bacterial pathogens [41, 42]. We investigated the life-span of HIV-1 virus in the two mouse strains since the mouse is not a normal permissive host. We tested the concentration and time of HIV-1 VL in the peripheral blood of the non-humanized mouse as a parameter that potentially could influence viral pathogenicity. On the first and second day after intraperitoneally injection of 0.3 ml of HIV-1_ADA_ viral stock, the detected copy numbers of HIV RNA were lower by ∼0.3 log_10_ in NSG-*cmah*^-/-^ mice compared to NSG [4.98 ± 0.1 and 4.1. ±0.08 compared to 5.25. ± 0.06 and 4.4 ± 0.05 log_10_, respectively (P < 0.05, **Fig. 8a**)].

**Fig. 8.**
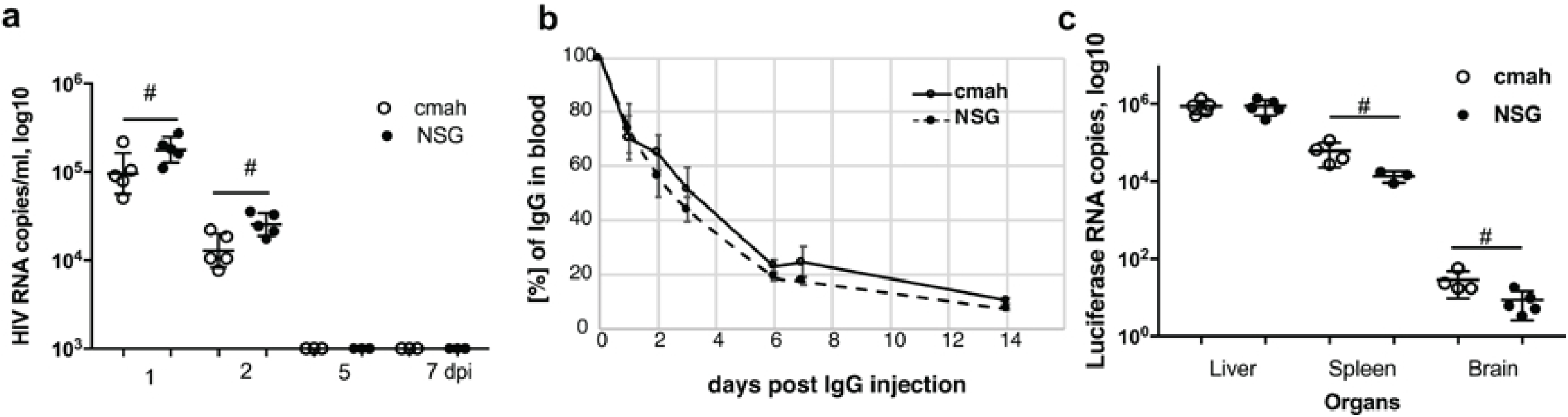
Effects of *cmah*^-/-^ background on HIV-1 and IgG clearance from blood circulation and tissue luciferase expression delivered by rAAV2/DJ8. **a** Clearance of HIV-1 from mouse peripheral blood. **b** Clearance of human IgG from the peripheral blood (n = 5). Percentage of human IgG determined at 30 minuntes after injection of 100 μl of a 10% human IgG (10 mg/mouse) were 10.6 ± 1.2 and 11.0 ± 0.9 μg/ml for NSG-*cmah*^-/-^ and wild type mice, respectively. Five to three blood samples were collected for days 1 – 14 time points, mean and SEM are shown. End-point IgG concentrations in peripheral blood were 1.1 ± 0.04 and 0.9 ± 0.03 μg/ml for NSG-*cmah*^-/-^ and wild type mice, respectively. **c** Luciferase RNA expression at 32 days post intravenous injection in liver, spleen, and brain. (#) P ≤ 0.05 were considered significant by one-tail Mann-Whitney test.

The clearance of sialylated glycoproteins going through the multiple types of receptors and changes of mouse sialylation could affect interaction with human immune globulins. We compared the time of circulation of human IgG in NSG-*cmah*^-/-^ and wt mice (**Fig. 8b**). We were not able to determine the first minutes of injected IgG dose clearance, but overall clearance of human IgG was nearly identical in the two strains of mice. A slightly higher concentration remained in NSG-*cmah*^-/-^ mice compared to NSG at the end of observation (1.1 ± 0.04 and 0.9 ±0.03 μg/ml for NSG-*cmah*^-/-^ and wt mice, respectively) (P < 0.05).

Biodistribution and expression levels of recombinant adeno-associated virus (rAAV) vector delivered genes could be affected by sialylation of host receptors [5, 10, 43]. We compared the luciferase expression delivered by rAAV2/DJ8. We were not able to detect statistically significant differences in luminescence in liver and spleen between the two strains at 7, 14, and 21 days post-inoculation (data not shown). However, at 32 days the expression of luciferase RNA was significantly higher in the spleen and brain of NSG-*cmah*^-/-^ as determined by ddPCR (**Fig. 8c**). It was higher by 0.60 log_10_ copies in spleen (4.72 ± 0.12 vs 4.12. ± 0.07 log_10_) and 0.54 log_10_ in brain (1.40 ± 0.11 vs 0.85. ± 0.12 log_10_) of *cmah*^-/-^ mice compared to NSG (P < 0.05).

## Discussion

We created NSG-*cmah*^-/-^ mice with the intent to use this strain for different aspects of biomedical research. As initial phenotypic characterization, we compared several parameters involved in human immune system development in NSG-*cmah*^-/-^ to the parent NSG strain. Glycoproteins-lectin interactions (for example, hematopoetic cells surface glycoporteins – bone marrow stromal cell lectins/selectins) are an important mechanism of human B-cell selection. The NSG mouse environment that contained Neu5Gc (non-human type of sialylation) exhibited a higher frequency of clonal B-cell outgrowth that may represent responses against the Neu5Gc moiety. In the absence of Neu5Gc, human B cells appear to remain less activated. This finding indicates that non-human sialylation has a negative effect on human B cell development, and the NSG-*cmah*^-/-^strain provides a more supportive environment with good repertoire development and less clonal outgrowth [44]. The observations of lower IgD+ and CD22+ human B cells and sustained naïve CD45RA+CD3+ cells in the NSG-*cmah*^-/-^ strain also support this conclusion.

We observed better bone marrow B-cell precursor engraftment in control NSG mice compared to NSG-*cmah*^-/-^, and better bone marrow T-cell engraftment in NSG-*cmah*^-/-^. The HIV-1 infected NSG-*cmah*^-/-^ mice showed a significant reduction in bone marrow T-cells and a subsequent increase in bone marrow B-cell precursor engraftment. These findings may indicate T-cell suppression of B-cell engraftment is occurring in this mouse strain but will require further study to confirm. In contrast, the NSG strain showed similar levels of bone marrow T-cell and B-cell engraftment regardless of infection with HIV-1.

The T cell TCRβ chain repertoires in NSG-*cmah*^-/-^ and wt NSG mice were not different in clonality metrics and remain polyclonal in bone marrow and spleen. However, compared to wt NSG mice, NSG-*cmah*^-/-^ exhibited better evenness of repertoire between spleen and bone marrow. We did not observe statistically significant differences in clonal T-cell expansion based on TCRβ CDR3 length frequency in bone marrow versus spleen. This was previously reported for NOD/scid mice transplanted with cord blood derived hematopoietic CD34+ stem cells [45].

In the studies presented here, we are not showing effects of vaccination with the childhood vaccines DTaP, HiB, and MMRII, which were tested in comparison on both strains of humanized mice at 6 months of age. Only a few animals developed antigen-specific IgM. The inability of CD34-NSG humanized mice to efficiently respond to vaccination is related to multiple deficiencies recognized in the humanized animals. This includes the absence of supportive cytokines, human follicular dendritic cells responsible for accumulation of antigen and stimulation of germinal center B-cells, deficiency of follicular helper cells involved in T-cell dependent immune responses, deficiency of complement, and others (reviewed in [46]). Several approaches to improve human adaptive immune responses were explored by elimination of mouse tissue histocompatibility antigens and introducing human MHC class I and II, cytokines and co-transplantation of bone, spleen, liver, and thymus. These approaches improved adaptive responses to varying degrees. It is possible that the introduction of these additional factors on the NSG-*cmah*^-/-^ background will further optimize the function of the human immune system and development of adaptive immune responses.

In addition to the comparison of human immune system cell development and phenotype, we assessed the behavior of human mature peripheral blood mononuclear cells in their ability to colonize mouse spleen and peripheral blood. In contrast to lymphocytes generated from human stem cells transplanted into NSG-*cmah*^-/-^ mice, which showed reduced levels of activation, transplanted mature lymphocytes very quickly expand and lose the naïve CD45RA+ phenotype. This could be attributed to the selection of human T cells in mouse thymus and reduced responsiveness to the mouse MHC (major histocompatibility) antigens.

Sialic acid on cellular membrane molecules has been identified as an attachment receptor for several pathogens and toxins. The composition could influence the viral capture by different cell types and *trans*-infection [47]. We observed increased HIV-1 mediated depletion of human CD4+ T-cells in the NSG-*cmah*^-/-^ strain of mice compared to the parental NSG. We also found a reduction of circulating HIV-1 RNA in non-engrafted NSG-*cmah*^-/-^ mice suggesting the viral particles may be more efficiently absorbed by cells with human-like sialylation patterns. We previously showed that mice transplanted with human hepatocytes also had enhanced clearance of virus from the circulation [48]. The increased pathogenicity of HIV-1 in NSG-*cmah*^-/-^ mice may be related to both the properties of human cells raised in the more human-like modified mouse environment as well as interaction of virus with the modified mouse stroma.

As humanized mice serve as a model to test biologic activity of human therapeutic antibodies [49, 50], we compared the clearance of human IgG. Human IgG contain oligosaccharides with N-acetylneuraminic acid (Neu5Ac, NANA); whereas, rhesus, cow, sheep, goat, horse, and mouse IgGs contain oligosaccharides with N-glycolylneuraminic acid (Neu5Gc, NGNA). The asialoglycoprotein receptor on hepatocytes [51] and FcRn on endothelial cells and placenta [52] could potentially be affected by human-like sialylation of receptors. Human FcRn binds to human, rabbit, and guinea pig IgG, but not significantly to rat, bovine, sheep, or mouse IgG (with the exception of weak binding to mouse IgG2b). In contrast, mouse FcRn binds to all IgG as previously analyzed [52]. Our observation of slower clearance of human IgG injected intravenously in NSG-*cmah*^-/-^ mice suggests the model may be useful for evaluation of the therapeutic efficacy of human antibodies. Another miscellaneous application of NSG-*cmah*^-/-^ mice could be testing of the transduction efficacy of viral vectors as well as viruses [53]. We used a common vector of rAAV serotype 2 with luciferase and expected to see the differences of transduction efficacy between wt NSG and NSG-*cmah*^-/-^ mice. We found the expression of luciferase was not different in liver (the major affected organ), but we did observe differences in less commonly affected organs – such as spleen and brain. These findings suggest that endothelial and splenic hematopoietic cells with human-like sialylation profiles could be more sensitive to viral infection.

## Conclusions

Humanized mice are widely used to study the human immune system responses to pathogens and therapeutics. However, mouse specific glycosylation affects the development of the human immune system and responses to various agents – such as viruses or biological, human-specific products like antibodies. We demonstrated that human-specific sialylation established by mutation of the *CMAH* gene supports naïve B and T cell generation with polyclonal receptors repertoires. In contrast to NSG wild type mouse sialylation background, we found the NSG-*cmah*^-/-^background increased sensitivity to HIV-1 infection and influenced rAAV vector transduction patterns. As such, NSG-*cmah*^-/-^ mice may accelerate translational research that target human infections and therapeutics development.

## Material and Methods

### Animals

Strains obtained from the Jackson Laboratories C57Bl/6 (Stock No: 000664), CMAH knock-out (B6.129X1-*Cmah*^*tm1Avrk*^/J, Stock No: 017588), and NSG (Stock No:05557) were bred and housed in the pathogen-free animal facility at the University of Nebraska Medical Center (Omaha, NE, USA).

Generation of B6.129X1-*Cmah*^*tm1Avrk*^/J referred hereafter as C57Bl/6-Cmah^-/-^ was described previously [54]. In this mouse, exon 6 was deleted by targeting a cassette containing LoxP flanked exon 6 with a neomycin cassette into 129/SvJ derived ES cells, followed by removal of the exon and the neomycin cassette through Cre enzyme and injection of deleted clone into blastcysts to generate chimera. The mice line was backcrossed to C57BL6/J strain for 10 generations.

### CRISPR reagents, mice generation, and genotyping

We deleted exon 6 using CRISPR approach in NSG strain background. Two guide RNAs were identified to target exon 6 (Cmah gRNA 1 CTTTGTGCATTTAACGGACCTGG and Cmah gRNA 2 TGAAATATATCAACCCTCCAGGG). The sgRNAs were transcribed from DNA templates generated by annealing two primers using the HiScribe(tm) T7 Quick High Yield RNA Synthesis Kit (New England Biolabs, Ipswich, MA) following manufacturer’s instructions. Cas9 mRNA was prepared using the pBGK plasmid as described in [28]. Injection mixture was prepared by dilution of the components into injection buffer (5 mM Tris, 0.1 mM EDTA, pH 7.5) to obtain the following concentrations: 10 ng/μl Cas9 mRNA, 10 ng/μl Cmah Left Guide and Right Guide RNA. Female NSG mice 3–4 weeks of age (JAX Laboratories, Bar Harbor, ME, USA) were superovulated by intraperitoneal injection with 2.5IU pregnant mare serum gonadotropin (National Hormone & Peptide Program, NIDDK), followed 48 hours later by injection of 2.5 IU human chorionic gonadotropin (hCG, National Hormone & Peptide Program, NIDDK). Mouse zygotes were obtained by mating NSG stud males with superovulated NSG females. The animals were sacrificed 14 hours following hcG administration, and oviducts were collected to isolate fertilized embryos. Injection mixture was introduced into the pronuclei and cytoplasm of fertilized NSG embryos by microinjection using a continuous flow injection mode. Surviving embryos were surgically implanted into the oviducts of pseudopregnant CD-1 recipient females. Genomic DNAs were extracted from tail biopsies and genotyping PCRs were performed as previously reported [28]. Cmah Forward TCCCAGACCAGGAGGAGTTA and Cmah Reverse TCCACTCCGAGTTTCAGATCA primers were used for screening by PCR. The expected amplicon sizes were 455 bases and 428 bases respectively for wild type and mutant mice. The PCR products were column purified and were sequenced using the primers used for PCR amplification.

### Western blot and flow cytometry analyses of NeuGc expression

Tissue samples from NSG and NSG-*cmah*^-/-^ mice were collected and homogenized in ice cold RIPA buffer with HALT protease inhibitor (cat#78430, ThermoFisher Scientific, Waltham, MA, USA). 20 μg of protein/sample was denatured in Laemmli sample buffer, then loaded and ran on a 12% SDS-polyacrylamide gel. The separated proteins were transferred to polyvinylidene difluoride (PVDF) membrane, blocked in 0.5% cold fish gelatin in TRIS buffered saline with 0.05% Tween-20 (TBS-T) for 2 hours at room temperature and then incubated in the primary antibody, Neu5GC (1:4000 in 0.5% cold fish gelatin TBS-T; chicken polyclonal IgY antibody, Biolegend Cat. No 146901) for overnight at 4 °C. After washes, the membrane was incubated in HRP-conjugated goat-anti-chicken IgY (GeneTex Cat. GTX26877, Lucerna-Chem AG, Switzerland) secondary antibody (1:4000) for 1 hour at room temperature. The immunoblot was developed with chemiluminescence and imaged using FluorChem M imager (Proteinsimple, San Jose, CA, USA). GAPDH was used as housekeeping control (clone GA1R, cat #MA5-15738, Thermoscientific, Waltham, MA USA). For flow cytometry, whole blood was obtained by bleeding the mice from the facial vein. After spinning it down at 1800 rpm for 8 minutes and removing the excess serum, 50 μl of the 1:1 mixture of the whole blood and 0.5% gelatin from cold water fish was incubated in the Fc block for 15 minutes on ice. Next, the primary anti-Neu5Gc was added and incubated for 30 minutes on ice. Washing was done by re-suspending the samples in 1ml of the 0.5% gelatin from cold water fish and spinning it at 1500 rpm for 5 minutes for 3 times. The secondary FITC-labeled anti-chicken reagent (**ab46969**) staining followed by red blood cells lysis using red blood cell lysis (Cat. #349202 from BD Bioscience, San Jose, CA) were performed. Further, acquisition was done on BD LSR II flow cytometer cells and analyzed on gated leukocytes using FLOWJO, analysis software (Tree Star, Ashland, OR, USA).

### Immunohistology

Organs collected from NSG-*cmah*-/-, NSG wt and C57Bl/6 mice were fixed and paraffin embedded; 5 μ thick sections were stained with antibodies for Neu5Gc (anti-Neu5Gc antibody, Biolegend, San Diego, CA, USA, Cat. No: 146903, dilution 1:100) at 4 °C overnight. Secondary anti-chicken antibodies (Biolegend Cat. No. 146901) were used at dilution 1:100 and DAB as chromogen. Tissues counterstained with hematoxylin.

### Human PBMC transplantation and engraftment analyses

NSG-*cmah*^-/-^ and wild type NSG mice at 8 weeks of age were transplanted with single donor human PBMC intraperitoneally (20 × 10^6^ cells/mouse). Blood samples were collected by facial vein bleeding for immunophenotyping at variable intervals (7, 14, 24, 31 days). Five-week post-injection, mice were sacrificed; blood, liver and spleen tissues were collected for immunophenotyping and immunohistochemistry. Briefly, 50 µL of whole blood was stained in EDTA-coated tubes with two different monoclonal antibody panels (Table 1) in a final volume of approximately 100µl. Cells not stained with any of the antibody were initially used to define the gating strategy. Compensation beads (BD Biosciences, cat. #552843) were stained with each antibody separately and run at each acquisition to calculate the compensation matrix. Immunophenotyping was performed to determine the absolute count and frequency of blood cells: leukocytes (CD45, BD Biosciences, Cat. #555482); CD3 (#557943); CD4 (#560650); CD8 (#562428); CD19 (#562653) and CD14 (eBioscience, San Diego, CA, Cat. #17-0149-42) for blood and spleen, respectively. The gating strategy for identification of cell subsets is presented in Figure 1A. Presence of activation markers CD45RA/CD45RO (BD Biosciences, #560674; #563215) and CD62L (#555544) were also checked for CD4+ and CD8+ T cells subpopulations, CD22 on B cells (#563940) in blood and spleen as absolute count and frequency of parent, respectively. After staining of cells, red blood cells were lysed with FACS Lysing solution (BD Biosciences), and cells were washed with PBS and resuspended in 1% paraformaldehyde. For absolute counting of the blood samples, CountBrightTM absolute counting beads (Invitrogen, Carlsbad, CA, USA; catalog C36950) were added to each sample before acquisition. Acquisition of stained cells was performed on BD LSR II (BD Biosciences) using acquisition software FACS Diva (BD Biosciences); and data were analyzed using FLOWJO, analysis software (Tree Star). Event counts of each cell population were exported, and absolute count/µl of blood were calculated using the following formula: [(Number of cell events / number of bead events) × (assigned bead count of the lot (beads/50 µl) / Volume of sample)].

### Human CD34+ cell transplantation and engraftment analyses

NSG-*cmah*^-/^- and wild type NSG mice reconstituted with the same cord blood sample derived CD34+ cells (5×10^4^/mouse intrahepatically) were bled at three and again at six months of age. For both time points ∼100 μl of blood was collected into MiniCollect 0.5 mL EDTA tubes (Greiner Vacuette North America Inc., Monroe, NC, USA; Cat. #450475). Plasma was collected and stored at −80° for future use. Remaining cell samples were diluted at a 1:1 ratio with 50 μl FACS buffer (2% fetal bovine serum in PBS) and mixed thoroughly. The samples were divided into two panels, B cell and T cell. The B cell panel consisted of mouse anti-human CD45-FITC (BD Pharmingen; Cat. #555482), CD19-BV605 (BD Biosciences; Cat. #562653), CD22 BV421(#563940), IgG-PerCP/Cy5.5 (Biolegends, # 409312), IgD-PE (eBiosciences, #12-9868-42), IgM-APC (eBiosciences, #17-9998-42), and Brilliant Stain Buffer (BD Horizon, #563794) cocktails. The T cell panel consisted of: anti-human CD45-FITC (BD Pharmigen, #555482), CD3-Alexa Fluor 700 (BD Biosciences #557943), CD4-APC (BD Pharmingen #555349), CD8-BV421 (BD Horizons #562428), CD45RA-APC-H7 (BD Biosciences #560674), CD14-PE (BD Pharmingen #555398), and Brilliant Stain Buffer. After 30 minutes incubation at 4 °C, red blood cell lysis (BD Bioscience #349202) and two washes, samples were fixed with 2% paraformaldehyde and acquisition was performed on BD LSR II flow cytometer.

### HIV-1 infection

Animals with comparable peripheral blood repopulation with human leucocytes were infected with HIV-1_ADA_ at 10^4^ 50% tissue culture infectious (TCID_50_) dose intraperitoneally. At four and nine weeks post infection, animals were euthanized for the VL analysis (COBAS^®^ AmpliPrep/COBAS^®^ TaqMan^®^ HIV-1 Test, v2.0, Roche Molecular Systems Inc., Pleasanton, CA, USA) and human cells phenotypes by FACS. Peripheral blood, spleen, and bone marrow cells were analyzed as described above.

Additional bone marrow human population analysis was performed by flow cytometry at the end of observation at 5-6 weeks post infection. Approximately 10^6^ isolated bone marrow cells were aliquoted into three tubes for each mouse evaluated. The cells were washed once with 2 mL phosphate buffered saline (PBS) and were then incubated with the 8 antibody cocktails for 30 minutes at room temperature in 200 uL of PBS with 10% fetal calf serum (PBS-FCS). The following antibodies were used in combination: T-NK cocktail consisting of APC-H7 conjugated anti-CD3 (clone SK7), PE-Cy7 conjugated anti-CD4 (clone SK3), PE conjugated anti-CD7 (clone 8G12), PerCP-CY5-5 conjugated anti-CD8 (clone SK1), FITC conjugated anti-CD14 (clone MLP9), V450 conjugated anti-CD16 (clone 3G8) APC conjugated anti-CD56 (clone NCAM16.2) and V500c–conjugated CD45 (clone 2D1); MYELO cocktail consisting of APC-H7 conjugated anti-HLA-DR (clone L243), PE conjugated anti-CD10 (clone H10a), PE-Cy7 conjugated anti-CD13 (clone L138), APC conjugated anti-CD24 (clone 2G5), V450 conjugated anti-CD33 (clone WM53), FITC conjugated anti-CD34 (clone 8G12), PerCP-CY5-5 conjugated anti-CD117 (clone 95C3) and V500c–conjugated CD45 (clone 2D1) and the BCL tube consisting of FITC conjugated anti-kappa (clone TB28-2), PE conjugated anti-lambda (clone 1-155-2), PE-Cy7 conjugated anti-CD10, PerCP-Cy5.5 conjugated anti-CD19 (clone SJ25C1), APC-H7 conjugated anti-CD20 (clone L27), APC conjugated anti-CD24 (clone 2G5), V450 conjugated anti-CD38 (clone HB7) and V500c–conjugated CD45 (clone 2D1). All antibodies were obtained from BD Biosciences (Franklin Lakes, NJ, USA) except CD24 and CD117, which were obtained from Beckman Coulter (Brea, CA, USA).

After incubation, the cells were washed twice with 1 mL PBS and were resuspended in 500 uL PBS containing stabilizer (BD Biosciences) to fix the cells. Fifty thousand to 100,000 cell events were collected for each sample on a Becton Dickinson FACS Canto II (Franklin Lakes, NJ, USA). Analysis of the FCS files was performed using Kaluza 1.3 analysis software (Beckman Coulter).

Gating schemes for the bone marrow analysis are shown in the additional file 6 **Fig. S6**.

For all bone marrow cell populations characterized, total cell events were derived based on gating and logical antigen (Boolean) profiles. Population frequencies were then derived by dividing the specific cell population events by the total human cell events after CD45 and singlet gating.

### HIV-1 viral clearance

Viral clearance naïve non-humanized animals were injected with HIV-1 stock 3×10^3^ TCID_50_/mouse), and blood was collected on days 1, 2, 5, and 7 post inoculation. Number of viral RNA copies were determined as described above.

### Recombinant adeno-associated virus biodistribution

NSG-*cmah*^*-/-*^ and wild type NSG mice were injected intravenously with rAAV2/DJ EF1a-luciferase 1.825E+11 GE/mouse (Viral Vector Core, University of Iowa, Iowa City, IA; Cat No. Uiowa-6161: Lot AAV3240; http://www.medicine.uiowa.edu/vectorcore/). D-Luciferin Bioluminescent Substrate (Cat 770504, PerkinElmer, Waltham, MA) was used for in vivo detection. The biodistribution of luminescence were analyzed by IVIS^®^ Spectrum an in vivo imaging system at 1, 2, and 4 weeks after rAAV administration to validate and compare transduction efficiency. Liver, spleen, and brain tissues were collected at day 32 for detection of luciferase gene on droplet digital PCR (ddPCR) (QX200(tm) Droplet Digital(tm) PCR System, Bio-Rad, Hercules, CA, USA) with forward primer 5’CTTCGAGGAGGAGCTATTCTT-3’, reverse primer 5’-GTCGTACTTGATGAGAGTG-3’, and luciferase probe 5’-/56 FAM/TGCTGGTGC/ZEN/CCACACTATTTAGCT/3IABKFQ/-3’ (Integrated DNA Technologies, Inc., Coralville, IAUSA). Briefly, isolation of RNA was performed using a RNeasy Plus Universal Kit (Qiagen, Hilden, Germany) as per the manufacturer’s instructions. The final PCR reaction was comprised of ddPCR supermix (Bio-Rad), 20U/μl reverse transcriptase, 15mM Dithiothreitol (DTT), 900nM primers, 250nM probe and 100ng of RNA template in a final volume of 20 μl and loaded into an eight-channel disposable droplet generator cartridge (Bio-Rad). Generated droplets were then transferred into a 96-well PCR plate, heat-sealed with foil and then amplified to endpoint using a BioRad C1000 Touch PCR cycler at 95 °C for 10 minutes then 40 cycles of 94 °C for 15 seconds and 60 °C for 1 minute (2 °C/s ramp rate) with a final step at 98 °C for 10 minutes and 4 °C hold. Plates containing amplified droplets were loaded into a QX200 droplet reader (Bio-Rad). Discrimination between negative droplets (no luciferase copies) and positives (with luciferase copies) was used to estimate concentration of targets (luciferase copies/ul) using QuantaSoft analysis software (Bio-Rad). The resulted copies were normalized to input RNA and represented as luciferase copies in log scale.

### Human IgG clearance

We compared the circulation time of human IgG (IVIG, PRIVIGEN, CSL Behring LLC). NSG-*cmah*^-/-^ and control mice were injected with 100 μl of 10% IVIG in saline. Blood samples were collected at 30 minutes after IVIG injection as point day 0. The actual concentration of human IgG in peripheral blood was measured at days 1, 2, 3, 6, 7, and 14. Plasma human IgG concentration was determined by ELISA kit (Bethyl Laboratories, Inc. Montgomery, TX, USA, cat# E80-104).

## List of abbreviations

CMAH: CMP-N-acetylneuraminic acid hydroxylase
HRP: Horseradish peroxidase
HSPC: Hematopoietic stem/progenitor cells
HIV-1: Human immunodeficiency virus type 1
IHC: Immunohistochemistry
Neu5Ac: N-acetylneuraminic acid
Neu5Gc: N-glycolylneuraminic acid
NSG: NOD/scid-IL2Rγ_c_^−/−^ mice
PBMC: Peripheral blood immune cells
rAAV: Recombinant adeno-associated viral vector
Siglecs: Sialic acid–binding Ig-like lectins
WB: Western blot
WT mice: Wild-Type mice

## Declarations

### Acknowledgements

We would like to thank Mr. Edward Makarov and Mr. Weimin Wang for technical assistance with animal work; the Elutriation Core Facility in the Department of Pharmacology and Experimental Neuroscience provided peripheral blood mononuclear cells; and the Flow Cytometry Research Facility, the DNA Sequencing Core Facility and the Comparative Medicine Department; Robin Taylor for editing assistance. All located at University of Nebraska Medical Center.

## Availability of data and materials

All data generated or analyzed during this study are included in this published article and its additional files.

## Authors’ contributions

SG and LYP conceived and designed the study. RSD, ABW, and YC performed in vivo studies on mice, flow cytometry. RSD, SM, and PSJ performed immunohistochemistry, qPCR, and Western blot procedures. CBG, RMQ, and DWH performed embryo isolation, microinjection, and generation of founder mice. ABW, SM, YC, and SMM performed cord blood collection, CD34+ cells isolation, FACS analysis of peripheral blood. SJP performed flow cytometry of bone marrow. RSD, SJP, CBG, SG, and LYP interpreted the results and wrote the manuscript. All authors have read and given approval of the final version of the manuscript.

## Ethics approval and consent to participate

All animal experiments were conducted following *NIH guidelines for housing and care of laboratory animals* and in accordance with the University of Nebraska Medical Center protocols approved by the institution’s Institutional Animal Care and Use Committee (protocol numbers 13-009, 14-100 and 06-071) in animal facilities accredited by the Association for Assessment and Accreditation of Laboratory Animal Care International.

## Consent for publication

Not applicable.

## Additional files

Additional file 1. **Fig. S1.** Human immune cells IGH genes CDR3 length frequency distribution in bone marrow. **a** Histograms of IgH CDR3 length in bone marrow of NSG-*cmah*^-/-^ and **b** NSG mice. **c** The NSG-*cmah*^-/-^ background was associated with a lower clonal B-cell outgrowth, and subsequently a lower maximal frequency of templates. **d** The frequency of CDR3 lengths can be seen to show a more gaussian distribution in the NSG-*cmah*^-/-^ mice although and the number of total IGH templates was not statistically different **e** Statistical analysis was performed immunoSEQ Analyzer (https://www.adaptivebiotech.com/). Five animals per strain were used.

Additional file 2. **Fig. S2.** IgH CDR3 length in bone marrow and spleen in NSG-*cmah*-/-(m158, 103, 96, 159, 154) and NSG wild type mice (M376, 3578, 3571, 3577, 3573). Arrows indicate shortening of CDR3 length in spleen (orange bars) compared to the bone marrow (blue bars) samples in the same animal.

Additional file 3. **Fig. S3.** IGH genes family usage.

**a** Individual IgH genes families usage in NSG-*cmah*^-/-^ and **b** NSG wt mice represent the summary of productive rearrangement frequency in spleen tissue samples. Similar profiles were found in bone marrow (not shown). **c** The frequency of V, D and J genes family usage in bone marrow and spleen tissue are highly similar in both strains as expected. As shown, there are no significant differences between NSG and NSG-*cmah*^-/-^ mice in overall IgH gene family usage. In both strains, a significant reduction in the frequency of IGHD02 usage was apparent in splenic B-cells compared to bone marrow. Mild increased frequencies in IGHD01, IGD06, and IGD07 usage was also noted in NSG-*cmah*^-/-^ splenic tissue. In NSG wt mice, only IGD03 usage was increased in splenic B-cells compared to bone marrow. For J family usage, an increase in IGHJ03 frequency was found in NSG-*cmah*^-/-^ splenic B-cells with a reduction of IGHJ05 and IGHJ06 usage. Statistical analysis was done by two-way ANOVA with Fisher’s LSD test. P <0.05 (*) considered significant.

Additional file 4. **Fig. S4.** Spleen TCRβ chain gene usage.

**a** Individual TCRβ chain gene families usage in NSG-*cmah*^-/-^ and **b** NSG wt mice represent the frequency of productive rearrangements TCRβ in spleen tissue samples. Similar profiles were found in bone marrow (not shown).

Additional file 5. **Fig. S5.** Human TCRβ chain genes CDR3 size frequency distributions in spleen and repertoire characteristic in bone marrow and spleen. **a** Histograms of TCRβ chain gene CDR3 length in bone marrow of NSG-*cmah*^-/-^ and **b** NSG mice. **c** NSG-*cmah*^-/-^ background was associated with lower clonal frequencies in spleen (Spl) compared to bone marrow (BM), which was statistically significant by one-way paired t-test (P = 0.0381) for NSG-*cmah*^-/-^ mice.

Five animals per strain were used. **d** The richness of TCRβ chain gene repertoire in both strains was higher in spleen compared to bone marrow (*P < 0.05), while the number of total TCRβ chain gene templates was not statistically different between strains (not shown). **e** The Pielou’ evenness of repertoires in the bone marrow and splenic compartments was not significantly different in both strains; however, the evenness was higher in the spleens of NSG-*cmah*^-/-^ mice (one tail paired t-test, P = 0.0312). Red dashed line highlights higher evenness of TCRβ in NSG-*cmah*^-/-^ strain compared to wild type NSG by one-tail Mann Whitney test (P = 0.0446).

Additional file 6. **Fig. S6.** Humanized bone marrow gating strategies.

For all tubes, the human cells were isolated based on human CD45 expression. The CD45-positive cells were secondarily isolated using a singlet gate (forward angle peak height vs. forward angle area) to eliminate cell doublets and triplets. The percentage of lymphocytes was enumerated based on two CD45 by light scatter displays so that cells had to be present in both gates to be considered lymphocytes. T-cells, NK-cells, and monocytes were enumerated using a T-cell/NK-cell cocktail containing CD3, CD4, CD7, CD8, CD14, CD16, and CD56. T-cells were identified as CD3-positive, low side light scatter events. The CD3-positive T-cells were further characterized for CD4 and CD8 expression to enumerate the helper and cytotoxic T-cell subsets. NK-cells were isolated using a low side light scatter (SS) gate on the CD45 by side light scatter histogram. The low SS cells were characterized for CD56 and CD16 expression to enumerate the two NK-cell subsets (not shown here). Monocytes were isolated using a Boolean logic gate as CD4^dim^ and not CD3-positive cells. Promonocytes and mature monocytes were further identified based on CD14 expression density. CD19-positive, CD10-positive and CD20 positive B-cells were enumerated using a B-cell specific cocktail containing kappa, lambda, CD10, CD19, CD20, CD24 and CD38. CD19-positive, low SS cells were gated to enumerate total B-cells and precursors. The CD19-positive B-cells were further characterized for expression of CD10 to identify the B-cell precursors and CD20 to identify transitional to mature B-cells. Myeloblasts, proB-cells, preB-cells, mast cells, and granulocytes were enumerated using a myeloid cell cocktail containing HLA-DR, CD10, CD13, CD24, CD33, CD34 and CD117. CD34-positive events were gated on a CD34 by SS histogram and were further characterized as myeloblasts or pro B-cells based on the CD45 by SS profile. Total B-cell precursors were isolated based on HLA-DR, CD10, and CD24 co-expression. PreB-cells were calculated using Boolean logic as total B-cell precursors and not proB-cells. Mast cells were enumerated as CD117^bright^ events on a CD34 by CD117 display. Finally, the granulocytes were estimated based on a CD45 by high SS gate that excluded the CD117^bright^ mast cells. Samples were run on a Becton Dickenson Canto II flow cytometer and analyzed offline using Kaluza software from Beckman Coulter.

Additional file 7. **Fig. S7.** Profile of human cells in NSG-*cmah*^-/-^ at 9 weeks post HIV-1 infection. NSG-*cmah*^-/-^ mice were infected with HIV-1_ADA_ intraperitoneally at 5 months of age. At 9 weeks post-infection, samples were collected for FACS analyses of the peripheral blood, spleen, and bone marrow. **A**, Human cell profile in peripheral blood. FACS gating strategies used: human CD45/CD3/CD14; CD3/CD4/CD8; CD4/CD45RO. **B**, For spleen additional analysis included: human CD45/CD14/CD123/CD1c; and CD45/CD19. **C,** Bone marrow analyses was done for CD45/CD3/CD19 and lineage negative CD3^-^/CD19^-^ human CD34+ and CD33-positive cells. Individual mouse and means with SEM are shown. P values were determined with Mann-Whitney test. P ≤ 0.05 were considered significant. Reconstituted at variable levels, NSG-*cmah*^-/-^ mice showed high sensitivity to HIV-1 infection with significantly decreased numbers of human T-cells (predominantly helper T-cells) and CD4+CD45RO+ memory T-cells in peripheral blood and spleen. In contrast, CD19-positive mature B-cells (spleen) and B-cell precursors (bone marrow) were significantly increased in NSG-*cmah*^-/-^ mice following HIV exposure.

## References

1 Crocker PR, Feizi T: Carbohydrate recognition systems: functional triads in cell— cell interactions. Current Opinion in Structural Biology 1996, 6(5):679–691.

2 Varki A, Gagneux P: Multifarious roles of sialic acids in immunity. Annals of the New York Academy of Sciences 2012, 1253:16–36.

3 Mietzsch M, Broecker F, Reinhardt A, Seeberger PH, Heilbronn R: Differential adeno-associated virus serotype-specific interaction patterns with synthetic heparins and other glycans. Journal of virology 2014, 88(5):2991–3003.

4 Wu Z, Asokan A, Grieger JC, Govindasamy L, Agbandje-McKenna M, Samulski RJ: Single amino acid changes can influence titer, heparin binding, and tissue tropism in different adeno-associated virus serotypes. Journal of virology 2006, 80(22):11393–11397.

5 Rao L, Albright BH, Corriher T, Murlidharan G, Asokan A: 42. Differential Transduction Profiles of AAV Vectors in a Mouse Model of Human Glycosylation. Molecular Therapy 2015, 23:S18–S19.

6 Springer SA, Diaz SL, Gagneux P: Parallel evolution of a self-signal: humans and new world monkeys independently lost the cell surface sugar Neu5Gc. Immunogenetics 2014, 66(11):671–674.

7 Varki A: Colloquium paper: uniquely human evolution of sialic acid genetics and biology. Proceedings of the National Academy of Sciences of the United States of America 2010, 107 Suppl 2:8939–8946.

8 Mikulak J, Di Vito C, Zaghi E, Mavilio D: Host Immune Responses in HIV-1 Infection: The Emerging Pathogenic Role of Siglecs and Their Clinical Correlates. Frontiers in Immunology 2017, 8:314.

9 Huang LY, Patel A, Ng R, Miller EB, Halder S, McKenna R, Asokan A, Agbandje-McKenna M: Characterization of the Adeno-Associated Virus 1 and 6 Sialic Acid Binding Site. Journal of virology 2016, 90(11):5219–5230.

10 Lisowski L, Dane AP, Chu K, Zhang Y, Cunningham SC, Wilson EM, Nygaard S, Grompe M, Alexander IE, Kay MA: Selection and evaluation of clinically relevant AAV variants in a xenograft liver model. Nature 2014, 506(7488):382–386.

11 Dankwa S, Lim C, Bei AK, Jiang RH, Abshire JR, Patel SD, Goldberg JM, Moreno Y, Kono M, Niles JC et al: Ancient human sialic acid variant restricts an emerging zoonotic malaria parasite. Nat Commun 2016, 7:11187.

12 Smith H, Cole JA, Parsons NJ: The sialylation of gonococcal lipopolysaccharide by host factors: a major impact on pathogenicity. FEMS microbiology letters 1992, 100(1-3):287–292.

13 Takahashi T, Takano M, Kurebayashi Y, Masuda M, Kawagishi S, Takaguchi M, Yamanaka T, Minami A, Otsubo T, Ikeda K et al: N-Glycolylneuraminic Acid on Human Epithelial Cells Prevents Entry of Influenza A Viruses That Possess N-Glycolylneuraminic Acid Binding Ability. Journal of virology 2014, 88(15):8445–8456.

14 Ghaderi D, Zhang M, Hurtado-Ziola N, Varki A: Production platforms for biotherapeutic glycoproteins. Occurrence, impact, and challenges of non-human sialylation. Biotechnol Genet Eng Rev 2012, 28:147–175.

15 Tangvoranuntakul P, Gagneux P, Diaz S, Bardor M, Varki N, Varki A, Muchmore E: Human uptake and incorporation of an immunogenic nonhuman dietary sialic acid. Proceedings of the National Academy of Sciences of the United States of America 2003, 100(21):12045–12050.

16 Taylor CE, Cobb BA, Rittenhouse-Olson K, Paulson JC, Schreiber JR: Carbohydrate moieties as vaccine candidates: targeting the sweet spot in the immune response. Vaccine 2012, 30(30):4409–4413.

17 Cao H, Crocker PR: Evolution of CD33-related siglecs: regulating host immune functions and escaping pathogen exploitation? Immunology 2011, 132(1):18–26.

18 Ikehara Y, Ikehara SK, Paulson JC: Negative regulation of T cell receptor signaling by Siglec-7 (p70/AIRM) and Siglec-9. J Biol Chem 2004, 279(41):43117–43125.

19 Varchetta S, Brunetta E, Roberto A, Mikulak J, Hudspeth KL, Mondelli MU, Mavilio D: Engagement of Siglec-7 receptor induces a pro-inflammatory response selectively in monocytes. PLoS One 2012, 7(9):e45821.

20 Buchlis G, Mingozzi F, Soto PC, Pearce O, Hui DJ, Varki AP, High KA: Intrinsically Hyperactive and Hyperproliferative CD8+ T Cells In Cmah-/- Mice as a Model of Human Gene Transfer Responses. Blood 2010, 116(21):3773–3773.

21 Büll C, Collado-Camps E, Kers-Rebel ED, Heise T, Søndergaard JN, den Brok MH, Schulte BM, Boltje TJ, Adema GJ: Metabolic sialic acid blockade lowers the activation threshold of moDCs for TLR stimulation. Immunology And Cell Biology 2016, 95:408.

22 Naito Y, Takematsu H, Koyama S, Miyake S, Yamamoto H, Fujinawa R, Sugai M, Okuno Y, Tsujimoto G, Yamaji T et al: Germinal center marker GL7 probes activation-dependent repression of N-glycolylneuraminic acid, a sialic acid species involved in the negative modulation of B-cell activation. Molecular and cellular biology 2007, 27(8):3008–3022.

23 Jellusova J, Nitschke L: Regulation of B cell functions by the sialic acid-binding receptors siglec-G and CD22. Front Immunol 2011, 2:96.

24 Nystedt J, Anderson H, Hirvonen T, Impola U, Jaatinen T, Heiskanen A, Blomqvist M, Satomaa T, Natunen J, Saarinen J et al: Human CMP-N-acetylneuraminic acid hydroxylase is a novel stem cell marker linked to stem cell-specific mechanisms. Stem Cells 2010, 28(2):258–267.

25 Masse-Ranson G, Mouquet H, Di Santo JP: Humanized mouse models to study pathophysiology and treatment of HIV infection. Curr Opin HIV AIDS 2018, 13(2):143–151.

26 Paulson JC, Macauley MS, Kawasaki N: Siglecs as sensors of self in innate and adaptive immune responses. Annals of the New York Academy of Sciences 2012, 1253:37–48.

27 Cao L, Diedrich JK, Kulp DW, Pauthner M, He L, Park SR, Sok D, Su CY, Delahunty CM, Menis S et al: Global site-specific N-glycosylation analysis of HIV envelope glycoprotein. Nat Commun 2017, 8:14954.

28 Harms DW, Quadros RM, Seruggia D, Ohtsuka M, Takahashi G, Montoliu L, Gurumurthy CB: Mouse Genome Editing Using the CRISPR/Cas System. Curr Protoc Hum Genet 2014, 83:15 17 11–27.

29 Ito M, Hiramatsu H, Kobayashi K, Suzue K, Kawahata M, Hioki K, Ueyama Y, Koyanagi Y, Sugamura K, Tsuji K et al: NOD/SCID/gamma(c)(null) mouse: an excellent recipient mouse model for engraftment of human cells. Blood 2002, 100(9):3175–3182.

30 Ishikawa F, Yasukawa M, Lyons B, Yoshida S, Miyamoto T, Yoshimoto G, Watanabe T, Akashi K, Shultz LD, Harada M: Development of functional human blood and immune systems in NOD/SCID/IL2 receptor {gamma} chain(null) mice. Blood 2005, 106(5):1565–1573.

31 Giltiay NV, Shu GL, Shock A, Clark EA: Targeting CD22 with the monoclonal antibody epratuzumab modulates human B-cell maturation and cytokine production in response to Toll-like receptor 7 (TLR7) and B-cell receptor (BCR) signaling. Arthritis research & therapy 2017, 19(1):91.

32 Duty JA, Szodoray P, Zheng NY, Koelsch KA, Zhang Q, Swiatkowski M, Mathias M, Garman L, Helms C, Nakken B et al: Functional anergy in a subpopulation of naive B cells from healthy humans that express autoreactive immunoglobulin receptors. J Exp Med 2009, 206(1):139–151.

33 Kirchenbaum GA, St Clair JB, Detanico T, Aviszus K, Wysocki LJ: Functionally responsive self-reactive B cells of low affinity express reduced levels of surface IgM. Eur J Immunol 2014, 44(4):970–982.

34 Thome JJ, Grinshpun B, Kumar BV, Kubota M, Ohmura Y, Lerner H, Sempowski GD, Shen Y, Farber DL: Longterm maintenance of human naive T cells through in situ homeostasis in lymphoid tissue sites. Sci Immunol 2016, 1(6).

35 Espeli M, Rossi B, Mancini SJC, Roche P, Gauthier L, Schiff C: Initiation of pre-B cell receptor signaling: Common and distinctive features in human and mouse. Seminars in Immunology 2006, 18(1):56–66.

36 Zhuo Y, Bellis SL: Emerging Role of α2,6-Sialic Acid as a Negative Regulator of Galectin Binding and Function. The Journal of Biological Chemistry 2011, 286(8):5935–5941.

37 Vasta GR, Feng C, Gonzalez-Montalban N, Mancini J, Yang L, Abernathy K, Frost G, Palm C: Functions of galectins as ‘self/non-self’-recognition and effector factors. Pathogens and disease 2017, 75(5).

38 Poluektova LY, Garcia-Martinez, J.V., Koyanagi, Y., Manz, M.G., Tager, A.M.: Humanized Mice for HIV Research, 1 edn: Springer International Publishing AG.; 2014.

39 Pearson T, Greiner DL, Shultz LD: Humanized SCID mouse models for biomedical research. Current topics in microbiology and immunology 2008, 324:25–51.

40 Vassena L, Giuliani E, Koppensteiner H, Bolduan S, Schindler M, Doria M: HIV-1 Nef and Vpu Interfere with L-Selectin (CD62L) Cell Surface Expression To Inhibit Adhesion and Signaling in Infected CD4+ T Lymphocytes. Journal of virology 2015, 89(10):5687–5700.

41 Takahashi T, Takano M, Kurebayashi Y, Masuda M, Kawagishi S, Takaguchi M, Yamanaka T, Minami A, Otsubo T, Ikeda K et al: N-glycolylneuraminic acid on human epithelial cells prevents entry of influenza A viruses that possess N-glycolylneuraminic acid binding ability. Journal of virology 2014, 88(15):8445–8456.

42 Galili U: Natural anti-carbohydrate antibodies contributing to evolutionary survival of primates in viral epidemics? Glycobiology 2016, 26(11):1140–1150.

43 Asokan A, Hamra JB, Govindasamy L, Agbandje-McKenna M, Samulski RJ: Adeno-associated virus type 2 contains an integrin alpha5beta1 binding domain essential for viral cell entry. Journal of virology 2006, 80(18):8961–8969.

44 Larimore K, McCormick MW, Robins HS, Greenberg PD: Shaping of Human Germline IgH Repertoires Revealed by Deep Sequencing. The Journal of Immunology 2012, 189(6):3221–3230.

45 Lin C, Chen S, Yang L, Tan Y, Bai X, Li Y: Evaluation of TCR Vβ subfamily T cell expansion in NOD/SCID mice transplanted with human cord blood hematopoietic stem cells. Hematology 2007, 12(4):325–330.

46 Skelton JK, Ortega-Prieto AM, Dorner M: A Hitchhiker’s guide to humanized mice: new pathways to studying viral infections. Immunology 2018, 154(1):50–61.

47 Izquierdo-Useros N, Lorizate M, McLaren PJ, Telenti A, Kräusslich H-G, Martinez-Picado J: HIV-1 Capture and Transmission by Dendritic Cells: The Role of Viral Glycolipids and the Cellular Receptor Siglec-1. PLOS Pathogens 2014, 10(7):e1004146.

48 Dagur RS, Wang W, Cheng Y, Makarov E, Ganesan M, Suemizu H, Gebhart CL, Gorantla S, Osna N, Poluektova LY: Human hepatocyte depletion in the presence of HIV-1 infection in dual reconstituted humanized mice. Biol Open 2018, 7(2).

49 Gruell H, Klein F: Progress in HIV-1 antibody research using humanized mice. Curr Opin HIV AIDS 2017, 12(3):285–293.

50 Wang M, Yao LC, Cheng M, Cai D, Martinek J, Pan CX, Shi W, Ma AH, De Vere White RW, Airhart S et al: Humanized mice in studying efficacy and mechanisms of PD-1-targeted cancer immunotherapy. FASEB J 2018, 32(3):1537–1549.

51 Park EI, Manzella SM, Baenziger JU: Rapid clearance of sialylated glycoproteins by the asialoglycoprotein receptor. J Biol Chem 2003, 278(7):4597–4602.

52 Ober RJ, Radu CG, Ghetie V, Ward ES: Differences in promiscuity for antibody–FcRn interactions across species: implications for therapeutic antibodies. International Immunology 2001, 13(12):1551–1559.

53 Neu U, Bauer J, Stehle T: Viruses and Sialic Acids: Rules of Engagement. Current opinion in structural biology 2011, 21(5):610–618.

54 Hedlund M, Tangvoranuntakul P, Takematsu H, Long JM, Housley GD, Kozutsumi Y, Suzuki A, Wynshaw-Boris A, Ryan AF, Gallo RL et al: N-glycolylneuraminic acid deficiency in mice: implications for human biology and evolution. Molecular and cellular biology 2007, 27(12):4340–4346.

